# Who Infects Whom? Exploiting Bacterial Minicells for Targeted Virome Enrichment and Phage-Host Interaction Analysis through an Integrated Metagenomic Approach

**DOI:** 10.64898/2026.04.08.717211

**Authors:** Arezoo Pedramfar, Emma Ensenat, Natalie S. Allcock, Andrew D. Millard, Edouard E. Galyov

## Abstract

Linking bacteriophages (phages) to their hosts remains a fundamental challenge to understanding microbial ecology, viral evolution, and horizontal gene transfer. Although phages are the most abundant biological entities on Earth, the majority of them remain uncharacterized due to the lack of efficient host-linking approaches. Traditional methods, such as plaque assays, have significant limitations as they depend on visible lysis and therefore fail to detect phages that do not form plaques. Conversely, shotgun metagenomics can recover viral genomes directly from environmental samples; however, it cannot directly link phages to their bacterial hosts. In this study, we addressed this limitation by tackling the critical question of “who infects whom?” through the development of a novel, culture-independent approach that utilises an anucleate bacterial minicells-based platform to enrich for phages capable of infecting a target bacterial host. To validate our approach, purified *Escherichia coli* minicells were exposed to a concentrated viral fraction derived from sewage samples. Genomic DNA from phages that successfully infected and interacted with the *E. coli* minicells was isolated, amplified, and sequenced. Metagenomic analysis revealed a distinct *E. coli*-specific virome, including several putatively novel phage species and genera. This platform effectively bridges the gap between culture-dependent and metagenomic methods, providing a scalable, host-targeted tool for identifying phage-host pairs. Our approach also opens new opportunities for studying phage-host interaction networks in complex microbial ecosystems and enhances our ability to investigate viral diversity, host specificity, and the ecological roles of phages in natural environments.

## Main

Bacteriophages (phages) shape microbial communities and hold therapeutic promise, yet studying phage-host interactions remains complex [1]. Plaque assays detect only phages that produce visible plaques under laboratory conditions, a process constrained by host physiology and viral replication dynamics [2]. Culture-independent approaches, including shotgun metagenomics, have revealed extensive viral diversity but provided limited resolution of direct phage-host associations [3-5]. High-throughput chromosome conformation capture (Hi-C) improves host linkage through proximity ligation, yet linkage is influenced by restriction site density and crosslinking efficiency, and sequencing depth, which can introduce taxon-specific biases [6]. Recent studies, such as viral tagging, partially addressed these limitations but remain constrained by technical complexity, non-specific adsorption, false positives, and an inability to distinguish adsorption from productive infection [7-9].

We hypothesised that bacterial minicells can function as a biological platform for capturing host-specific phages and, in combination with metagenomics, enable deeper investigation of phage-host interactions. Bacterial minicells are small, anucleate cells that form when mutations misposition the division septum during bacterial cytokinesis [10, 11]. Minicells retain membranes, peptidoglycan, ribosomes, RNA, proteins and surface receptors identical to those of their parental cells. These properties make minicells attractive tools for studying selective phage interactions, as minicells combine the structural completeness of intact cells with the advantage of lacking host DNA contamination. Unlike viral tagging, minicell-based selection requires no modification of phage particles and enables natural, host-specific interaction [7-9].

To test the hypothesis that bacterial minicells can be used to capture host-specific phages, we first optimised the production of *E. coli* minicells and tested their specificity with known cognate and non-cognate phages. We then applied this approach to identify coliphages within an environmental sewage sample (Supplementary Methods).

Purified minicells were validated microscopically and by plating for Colony-Forming Units (CFU) to quantify the number of residual viable cells. No CFUs were detected, indicating that residual viable cells were below the limit of detection of the assay (Supplementary Fig. 1A-B). As a proof of principle, a mixture of cognate coliphage yaya_002 and the non-cognate *Staphylococcus capitis* phages AA0033_AM1 and AA0033_AM2 was interacted with *E. coli* minicells (Supplementary Table 1). Plaque assays demonstrated a 90% reduction in the recovery of the cognate phage yaya_002 with no significant change for *Staphylococcus capitis* phages (Supplementary Fig. 2A), suggesting only yaya_002 had absorbed to minicells and was confirmed by microscopy (Supplementary Fig. 2B-C). Sequencing further demonstrated 238-fold enrichment of yaya_002 and depletion of AA0033_AM1 (undetectable) and AA0033_AM2, with host genome contamination below 0.18 ± 0.012% (Table 1; Supplementary Table 2). Thus, demonstrating that minicells function as a biologically selective platform that distinguishes cognate from non-cognate phages.

**Table 1.** Comparison of control and post-incubation mean CPM values for a phage mixture containing the cognate coliphage yaya_002 and the non-cognate *S. capitis* phages AA0033_AM1 and AA0033_AM2, along with read mapping to the *E. coli* PB114 genome before (control) and after incubation of the phage mixture with *E. coli* PB114 minicells.

| Phage Name | Control (CPM) | Enriched- CPM (Mean ± SD) | Ratio (Enriched/Control) % |
| --- | --- | --- | --- |
| yaya_002 | 4.208E+03 | 1.00E+06 ± 5.33 | 2.38E+04 % |
| AA0033_AM1 | 3.5677E+04 | Not detected | Not detected |
| AA0033_AM2 | 9.6012E+05 | 3.80 ± 5.33 | 3.96E-04 % |

CPM = Counts Per Million.

Having established minicells-based enrichment works, we applied the approach to sewage samples (Supplementary Table 3). Again, host DNA contamination was low (<1.2%) (Supplementary Table 4). *E. coli* minicell-enriched viromes (MEPB1-3) exhibited reduced alpha diversity and increased beta diversity relative to total viromes (TVB1-3), indicating consistent selection for *E. coli*-associated phages (Supplementary Fig. 3; Supplementary Tables 5-8).

CLR analysis showed a distribution centered on the global mean (-2.6 × 10^-1^□; SD = 2.71), with enrichment thresholds defined at ±2SD (±5.42 log□units). A discrete subset of vOTUs exceeded the +2SD threshold (n = 225), indicating selective enrichment with minicells, whereas the remaining vOTUs showed depletion (Fig. 1A). The coliphage APPB2_1 (Supplementary Table 3 and 9) which was isolated from the same sewage sample as the viromes were constructed, exhibited ΔCLR values near the +2SD threshold, independently validating the cutoff as conservative, yet biologically meaningful.

**Figure 1.**
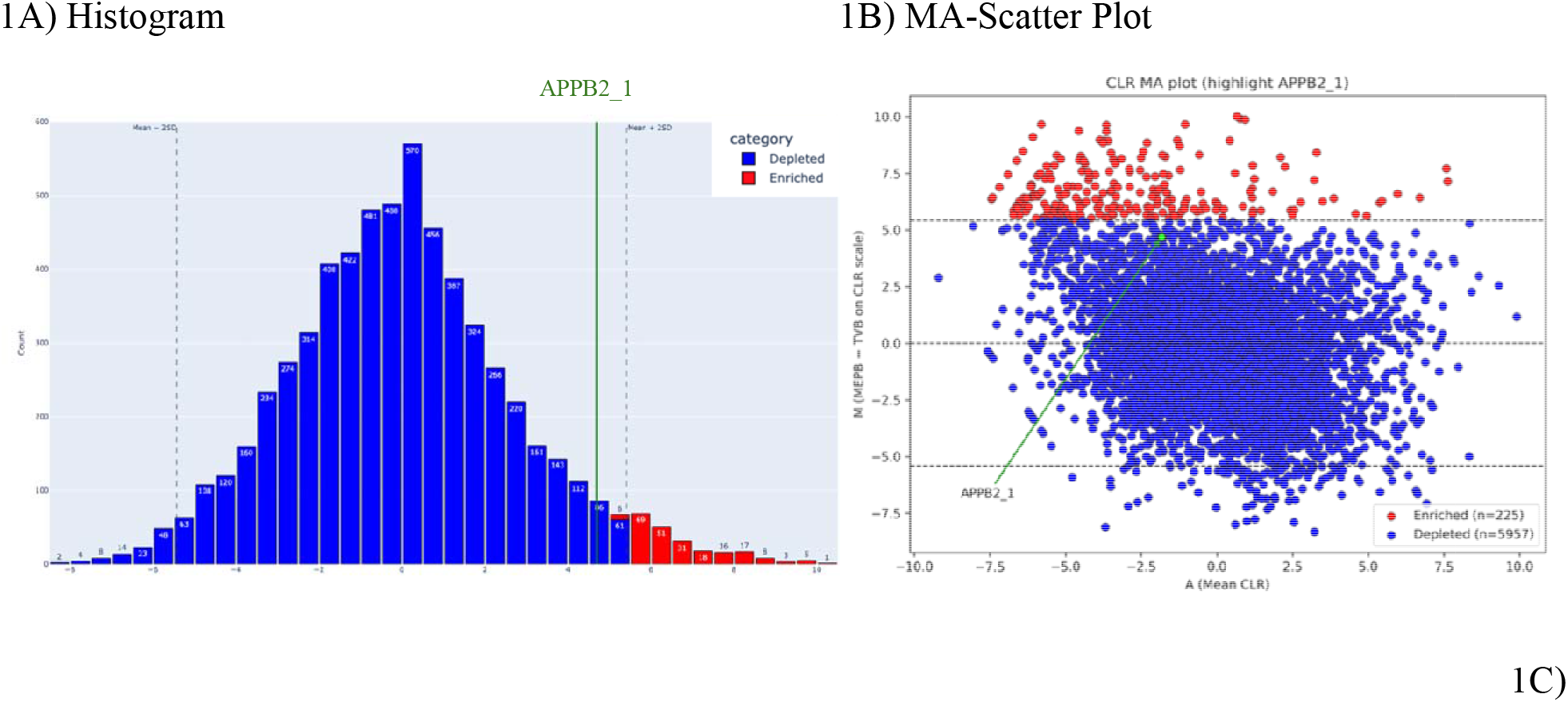
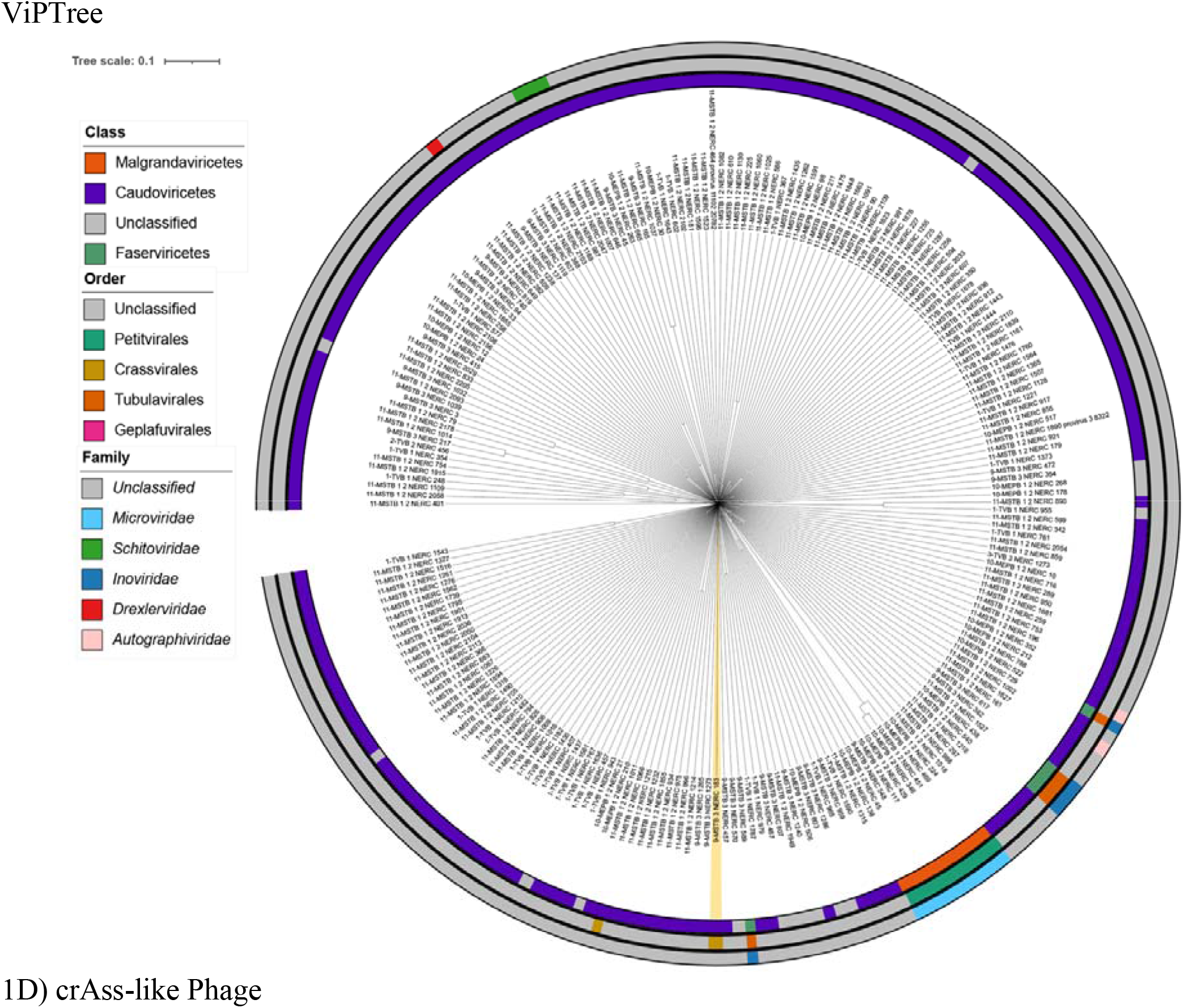
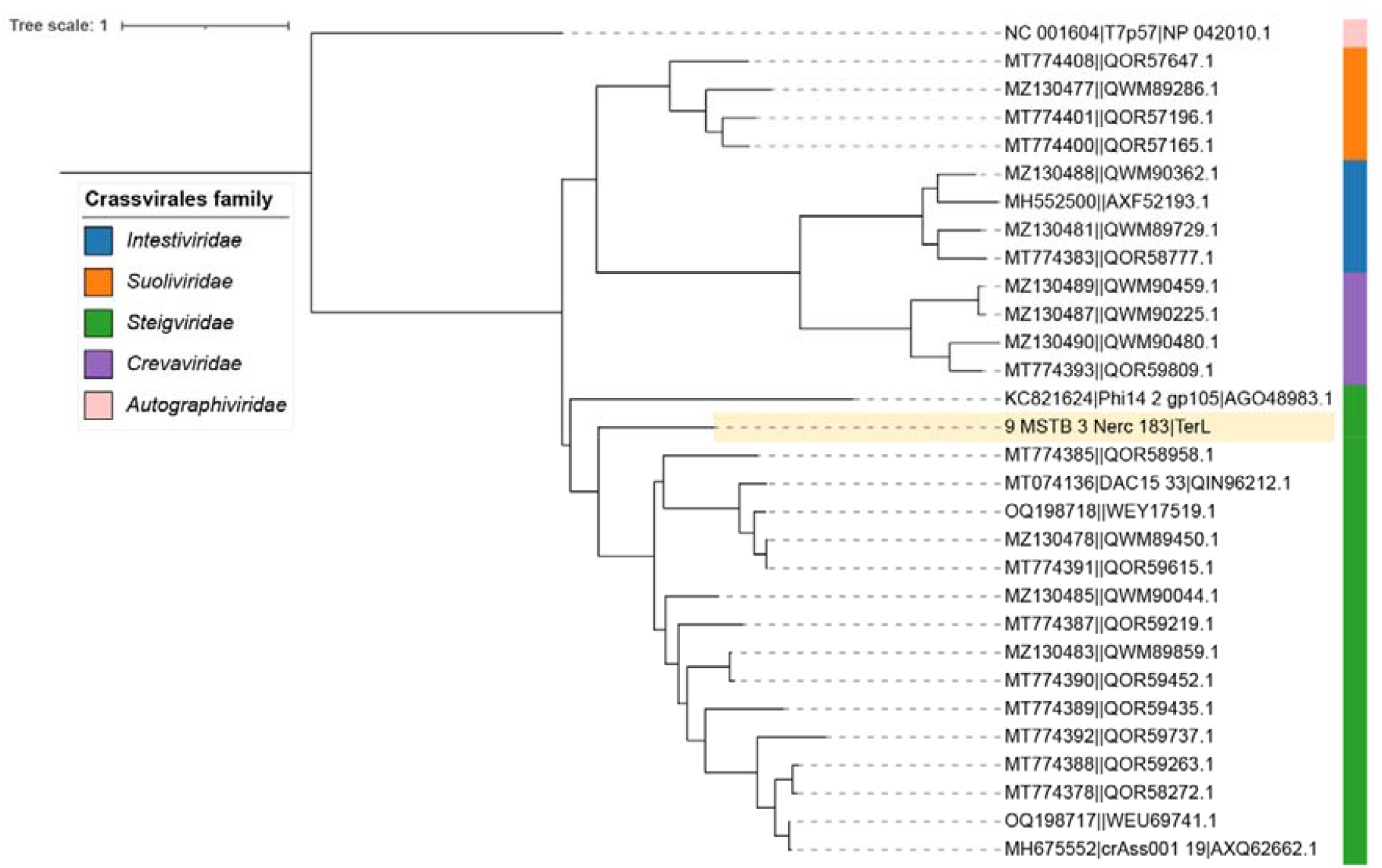
Selective enrichment and phylogenetic diversity of *E. coli* PB114-associated vOTUs. (1A) Histogram of ΔCLR (MEPB-TVB) values across vOTUs. Dashed lines indicate ±2SD thresholds (±5.42 log□ units). The right-hand tail represents enriched vOTUs following minicell-based selection. (1B) CLR mean-difference (MA) plot showed ΔCLR versus mean CLR abundance for each vOTU. Dashed lines mark ±2SD thresholds; red points denote enriched and blue points depleted vOTUs. The *E. coli* PB114-coliphage APPB2_1 is highlighted near the enrichment threshold. Data are derived from three biological replicates. (1C) Proteomic tree of 225 *E. coli* PB114-enriched vOTUs constructed using ViPTree and midpoint-rooted. The outer concentric rings indicate taxonomic assignments at class, order, and family levels. Most vOTUs were classified at the class level within *Caudoviricetes*, whereas taxonomic resolution decreased at the order and family ranks, with many genomes remained unclassified. (1D) Maximum-likelihood phylogeny inferred from TerL amino acid sequences of representative members of the order *Crassvirales*. The tree was rooted using the TerL sequence from phage T7 (NC_001604; NP_042010.1) as an outgroup from the family *Autographiviridae*. The TerL sequence encoded by vOTU 9-MSTB_3_NERC_183 (highlighted) clusters within the *Steigviridae* clade, close to Cellulophaga phage phi14:2 (KC821624). Branch lengths represent substitutions per site (scale bar shown).

The CLR-based MA plot (Fig. 1B) showed ΔCLR (MEPB–TVB) versus mean CLR abundance for each vOTU. vOTUs exceeding the +2 SD threshold were distributed across a broad range of mean abundances, consistent with selective enrichment rather than dominance by high-abundance taxa. The plaque-forming *E. coli* PB114 coliphage APPB2_1 lay close to the +2 SD cutoff, indicating that the selected threshold retained experimentally validated enriched vOTUs.

Of 6181 vOTUs recovered across total and enriched viromes, 26.4% indicated above 90% completeness (complete or high quality; Supplementary Table 10). Among 225 enriched vOTUs, 24.9% reached above 90% completeness (Supplementary Table 10). Host prediction using iPHoP (≥90% confidence) revealed increased representation of Enterobacterales-associated phages in minicells-enriched samples, alongside Bacteroidales and Pseudomonadales, and a reduced proportion of unclassified vOTUs relative to the total virome (Supplementary Fig. 5A-B).

Proteomic clustering (ViPTree) demonstrated that minicell-enriched vOTUs were distributed across multiple deep-branching clades, rather than forming a single cohesive cluster (Fig. 1C). Indicating that minicell enrichment did not preferentially amplify one closely related phage group but instead captured diverse members of the class *Caudoviricetes*. Genome-level Comparison with the INPHARED database (31 August 2025; 5 472 genomes) [12] revealed that 206 of 225 enriched vOTUs (91.6%) lacked detectable similarity to any sequenced reference genome, and only two exhibited moderate similarity (ANI ≈87-92%) (Supplementary Table 10-11). Thus, minicell enrichment expanded the diversity of putative coliphages, an already well-sampled group of phages compared to phages infecting most bacteria [13].

One of the most unexpected vOTUs detected in the minicells-enriched virome, 9-MSTB_3_NERC_183, corresponded to a crAss-like phage genome within the order *Crassvirales*. A gene encoding the large terminase (TerL) protein, commonly used as a phylogenetic marker for *Crassvirales* [14], was identified within this vOTU. Phylogenetic analysis placed this sequence within the family *Steigviridae*, a sister group to Cellulophaga phage phi14:2 (KC821624; Fig. 1D), and may represent the first *Crassvirales* phage associated with *E. coli* cells.

In this platform, minicells allow phage adsorption and DNA injection but not replication; fold changes reflect adsorption rather than phage replication. However, injection of phage DNA alone into minicells does not ensure productive infection in that host would occur [15].

A limitation of this approach is the dependency on genetically modified rod-shaped bacteria for minicell production, which constrains universal applicability. When genetically amenable bacteria are available, the ability to add or remove specific receptors to bacteria producing minicells will all selection of both host and receptors specific viromes in the future. In combination with advances in long-read sequencing technologies and cell sorting techniques, this would allow the reconstruction of complete genomes directly linked to host cells.

In conclusion, the minicell-based platform enables selective isolation of cognate phages from environmental phage mixtures while minimising host-genome contamination. This platform bridges the gap between culture-based and metagenomic approaches, providing a scalable method for viral discovery and host linkage and opening new avenues for exploring phage-host interactions across complex microbial ecosystems.

## Author Contributions

E.E.G conceived the study. A.P performed the experiments. E.E assisted with SEM/TEM sample preparation. N.S.A developed the SEM/TEM methodology and generated microscopy data. A.P, A.D.M and E.E.G analysed the data. A.P, A.D.M and E.E.G drafted the manuscript. A.D.M and E.E.G supervised the work. All authors reviewed and approved the final manuscript.

## Acknowledgment

This study, and A.P., were funded under project reference 2592104 by the Biotechnology and Biological Sciences Research Council (BBSRC) through the Midlands Integrative Biosciences Training Partnership (MIBTP) Doctoral Training Partnership (DTP), the Natural Environment Research Council (NERC; SSP 200580; NEOF/NBAF), and the pump priming from Leicester Microbial Sciences and Infectious Diseases Centre (LeMID). A.D.M was supported by MRC (MR/T030062/1). This work utilised the ALICE High Performance Computing (HPC) facility at the University of Leicester. The authors thank Professor Piet de Boer (Case Western Reserve University) for providing the *Escherichia coli* PB114 minicell-producing strain; Sayde Perry for providing the yaya_002 phage; and Abdullah A. A. Alahmadi for providing phages AA0033_AM1 and AA0033_AM2, as well as *Staphylococcus capitis* strain EB3. The authors also acknowledge the University of Leicester Core Biotechnology Services Electron Microscopy Facility for technical support.

## References

1. Hyman P. Are you my host? An overview of methods used to link bacteriophages with hosts. Viruses. 2025;17:65 10.3390/v17010065

2. Panteleev V, Kulbachinskiy A, Gelfenbein D. Evaluating phage lytic activity: From plaque assays to single-cell technologies. Front Microbiol. 2025;16:1659093 10.3389/fmicb.2025.1659093

3. Paez-Espino D, Eloe-Fadrosh EA, Pavlopoulos GA et al. Uncovering earth’s virome. Nature. 2016;536:425–30 10.1038/nature19094

4. Roux S, Hallam SJ, Woyke T et al. Viral dark matter and virus–host interactions resolved from publicly available microbial genomes. eLife. 2015;4:e08490 10.7554/eLife.08490

5. Emerson JB, Roux S, Brum JR et al. Host-linked soil viral ecology along a permafrost thaw gradient. Nat Microbiol. 2018;3:870–80 10.1038/s41564-018-0190-y

6. Bickhart DM, Watson M, Koren S et al. Assignment of virus and antimicrobial resistance genes to microbial hosts in a complex microbial community by combined long-read assembly and proximity ligation. Genome Biology. 2019;20 10.1186/s13059-019-1760-x

7. Džunková M, Low SJ, Daly JN et al. Defining the human gut host–phage network through single-cell viral tagging. Nature Microbiology. 2019;4:2192–203 10.1038/s41564-019-0526-2

8. Deng L, Ignacio-Espinoza JC, Gregory AC et al. Viral tagging reveals discrete populations in synechococcus viral genome sequence space. Nature. 2014;513:242– 45 10.1038/nature13459

9. de Jonge PA, von Meijenfeldt FAB, Costa AR et al. Adsorption sequencing as a rapid method to link environmental bacteriophages to hosts. iScience. 2020;23:101439 10.1016/j.isci.2020.101439

10. Frazer AC, Curtiss R, 3rd. Production, properties and utility of bacterial minicells. Curr Top Microbiol Immunol. 1975;69:1–84 10.1007/978-3-642-50112-8_1

11. Farley MM, Hu B, Margolin W et al. Minicells, back in fashion. Journal of Bacteriology. 2016;198:1186–95 10.1128/jb.00901-15

12. Cook R, Brown N, Redgwell T et al. Infrastructure for a phage reference database: Identification of large-scale biases in the current collection of cultured phage genomes. PHAGE. 2021;2:214–23 10.1089/phage.2021.0007

13. Hatfull GF. Dark matter of the biosphere: The amazing world of bacteriophage diversity. Journal of Virology. 2015;89:8107–10 10.1128/jvi.01340-15

14. de Jonge PA, van den Born BH, Zwinderman AH et al. Phylogeny and disease associations of a widespread and ancient intestinal bacteriophage lineage. Nat Commun. 2024;15:6346 10.1038/s41467-024-50777-0

15. Mäntynen S, Laanto E, Oksanen HM et al. Black box of phage–bacterium interactions: Exploring alternative phage infection strategies. Open Biology. 2021;11:210188 10.1098/rsob.210188

