## Supplementary information for "Who Infects Whom? Exploiting Bacterial Minicells for Targeted Virome Enrichment and Phage-Host Interaction Analysis through an Integrated Metagenomic Approach"

### Supplementary data

#### Methods

##### Media and buffers

Luria-Bertani (Oxoid™, Thermo Fisher Scientific) medium was used as the primary culture medium. LB broth, LB agar (1.5% (w/v)), and top agar (0.5% (w/v)) were prepared according to standard formulations and supplemented with 1 mM Calcium chloride ($CaCl_2$; Merck). Saline magnesium (SM) buffer (5.8 g NaCl $L^{-1}$, 2.0 g $MgSO_4$ $L^{-1}$, 50 mM Tris-HCl, pH 7.5) and phosphate-buffered saline (PBS; Merck) were prepared in distilled water. All media and buffers were sterilized by autoclaving at 121 °C prior to use.

##### Strain, phages and growth conditions

*Escherichia coli* PB114 (PB103, *ΔminCDE::aph*; minicell-producing; De Boer, 1989; obtained from Dr Edouard Galyov's laboratory collection, which was provided by Professor Piet De Boer) was used throughout this study. Cultures were grown in LB broth supplemented with 1 mM $CaCl_2$ at 37 °C with shaking (200 rpm) or on LB agar plates containing 1 mM $CaCl_2$ at 37 °C. The stationary growth phase was typically reached after about 18 h. Long-term storage was achieved using glycerol stocks (50% glycerol) at -80 °C, with short-term storage on LB agar plates at 4 °C.

The phages used in this study included AA0033_AM1 and AA0033_AM2, both specific to *Staphylococcus capitis* strain EB3 and obtained from Dr Edouard Galyov's laboratory collection and isolated by Abdullah A. A. Alahmadi, and phage yaya_002, which infects *Escherichia coli* MG1655 (originating from the BMCPR collection, isolated by Sayde Perry). Phages AA0033_AM1 and AA0033_AM2 were propagated and assayed on *S. capitis* EB3, while phage yaya_002 was propagated on *E. coli* PB114.

Supplementary Table 1- Overview of proof-of-principle phage genomes and metagenomic sequencing datasets. This table presented isolated phage genomes and metagenomic sequencing samples, including sample descriptions, incubation conditions with *Escherichia coli* PB114, genome lengths (base pairs) of assembled phages, and raw read counts for the sequencing datasets. Raw read counts correspond to paired-end sequencing reads (R1 files), with forward and reverse reads present in approximately equal numbers. Genome lengths were calculated from assembled FASTA sequences. Run and study accession numbers are provided for datasets deposited in the European Nucleotide Archive (ENA).

| Name | Description | Incubated with *E. coli* PB114 | Genome Length (bp) / Raw Reads Counts | Run Accession (ENA) | Study Accession (ENA) |
|---|---|---|---|---|---|
| **Phage yaya_002** | Host: *E. coli* MG1655; propagated: *E. coli* PB114 | No | 52133 | To be updated | To be updated |
| **Phage AA0033_AM1** | Host and propagated: *S. capitis EB3* | No | 17842 | To be updated | To be updated |
| **Phage AA0033_AM2** | Host and propagated: *S. capitis EB3* | No | 16125 | To be updated | To be updated |
| **APEPYARD_001_S173** | Mixture of phage yaya_002, AA0033_AM1, and Aa0033_AM2 Enriched virome Replicate 1 | Yes | 273899619 | To be updated | To be updated |
| **APEPYARD_002_S174** | Mixture of phage yaya_002, AA0033_AM1, and Aa0033_AM2 Enriched virome Replicate 2 | Yes | 299647441 | To be updated | To be updated |
| **APEPYARD_003_S175** | Mixture of phage yaya_002, AA0033_AM1, and Aa0033_AM2 Enriched virome Replicate 3 | Yes | 457100384 | To be updated | To be updated |
| **APYARD_005_S177** | Mixture of phage yaya_002, AA0033_AM1, and Aa0033_AM2, (Control) | No | 310456172 | To be updated | To be updated |
| **NC_MEPB_proof** | Only Minicells of *E. coli* PB114 (Negative control) | – | Failed | – | – |

#### Minicell isolation and purification (*E. coli* PB114)

A single *E. coli* PB114 colony was inoculated into 400 mL of LB supplemented with 1 mM $CaCl_2$ and incubated for 22 hours (h) at 37 °C with shaking (200 rpm). Culture density, minicell enrichment, and viable cell counts were monitored by Optical density at 600 nm ($OD_{600}$; Jenway spectrophotometer) and colony-forming units (CFU) enumeration. Phosphate-buffered saline (PBS; Merck) was added to a final concentration of 10% (v/v), and minicells were isolated by sequential differential centrifugation 2 500 × g for 10 min to pellet bacterial cells, followed by 13 000 × g for 10 min to pellet minicells (Hitachi High-Speed Refrigerated Centrifuge CR22N). Minicells were resuspended in LB supplemented with 1 mM $CaCl_2$ and incubated for 30 min at 37 °C; ceftriaxone (100 µg/mL) was added after 15 min to inhibit parental cell wall synthesis. The differential centrifugation scheme was repeated to remove remaining bacterial cells and re-pellet minicells. Minicells were washed in PBS, passed through a 0.7 µm pore-size filter (Target2 GMF, Fisher Scientific UK Ltd) to remove residual intact bacteria, concentrated to 2 mL using a 0.22 µm pore-sized (Sarstedt Ltd) inverted filtration unit, supplemented with chloramphenicol (100 µg/mL), and stored at 4 °C until use.

**Minicell quantification**

Total minicell counts were estimated by $A_{600}$ using the formula: minicells/mL = $A_{600} \times 5.0 \times 10^{10}$, integrating $OD_{600}$, LPS concentration, and relative size (minicell ~1/5 parental cell diameter) (Giacalone et al., 2006; Jivrajani et al., 2013).

**Phage lysate preparation**

Each phage was propagated on log-phase *E. coli* PB114 cells at a multiplicity of infection (MOI) of 0.1 and incubated until culture clearing or overnight. Lysates were clarified by centrifugation (4 000 × g, 10 min) and filtered through a 0.22 µm pore sized membrane to remove residual cells and debris. Purified lysates were stored for short-term at 4 °C in SM buffer.

**Minicells-phage interaction assay**

The mixtures contained 500 µL cognate phage yaya_002 at a titer of $10^3$ PFU/mL, 500 µL non-cognate phages AA0033_AM1 and AA0033_AM2 at a titre of $10^8$ PFU/mL, and 500 µL of concentrated *E. coli* PB114 minicells (~$10^9$ cells/mL). Prior to interaction, minicells were treated with DNase I (20 mg/mL) for 30 min at 37 °C. A control contained *E. coli* PB114 minicells (~$10^9$ cells/mL) with 500 µL of SM buffer. Mixtures were incubated for 30 min at 37 °C. Samples were washed and inverted through a 0.22 µm pore sized, 13 mm filter (VWR International Ltd) using 500 µL of 5 mM Magnesium sulfate ($MgSO_4$; Merck). The filtrate was assayed by plaque assay. The same method was applied to the interaction between *E. coli* PB114 minicells and the concentrated total virome.

**DNA extraction and Whole-Genome Amplification (WGA)**

Phase 1 (pre-extraction): Phage mixtures lacking minicell interactions were treated with DNase I for 30 min at 37 °C, then washed and inverted through a 0.22 µm pore sized (13 mm) filter (VWR) using 500 µL of 5 mM $MgSO_4$. Phase 2 (DNA extraction): Samples were incubated with lysozyme (250 µg) and proteinase K (20 µg) for 1 h at 60 °C, followed by 10 min at 90 °C. DNA was extracted by repeated phenol:chloroform:isoamyl alcohol (25:24:1) extraction until the interphase was clear, followed by chloroform extraction. The aqueous phase was treated with RNase I (20 mg/mL) for 15 min at 37 °C and re-extracted with chloroform. DNA was precipitated with 0.1 vol 3 M sodium acetate (pH 7.5) and ethanol at -20 °C overnight, pelleted by centrifugation (18 000 × g, 4 °C), washed twice with 70% ethanol, air-dried, and resuspended in 10-25 µL nuclease-free water. Extracted DNA was purified using the Zymo DNA Clean & Concentrator kit and amplified using the GenomiPhi V2 DNA Amplification Kit (illustra) according to the manufacturer's instructions. DNA quantity and quality were assessed using NanoDrop and Qubit.

**Library preparation and sequencing (proof-of-principle datasets)**

Sequencing libraries were prepared by SEQCENTER (USA) using a tagmentation- and PCR-based workflow with custom 10 bp unique dual indices, targeting ~280 bp insert sizes. Libraries were sequenced on an Illumina NovaSeq X Plus platform (paired-end 2 × 151 bp). Demultiplexing, quality control, and adapter trimming were performed using bcl-convert v4.2.4. Sequencing depth targeted ~200 Mbp per sample for dsDNA viruses (genomes <5 Mbp), corresponding to approximately 1.33 million reads.

**Bioinformatics (proof of principle datasets)**

Quality control: FastQC v0.11.9 (default settings) [1]. Trimming: trim_galore v0.6.7 (trim_galore --paired read1.fastq.gz read2.fastq.gz) [2]. Host mapping (% contamination and coverage): bbmap.sh v38.84 (minid=0.95) [3]. Read counts: SeqFu 1.20.0 (count) [4]. Host removal: minimap2 v2.24-r1122 with options -a -x asm5 [5]; SAMtools v1.13 (-f 4) to retain only unmapped reads to produce host-depleted FASTQ files. Mapping to known phage mixtures: bbmap.sh v38.84 (minid=0.95) [3] against a reference comprising the fasta sequences of the phages present prior to interaction. Reads were mapped to a reference database comprising the FASTA sequences of phages present prior to interaction. Abundance was first calculated as reads per kilobase per million mapped reads (RPKM), normalizing read counts to both phage genome length (kilobases) and total mapped reads ($\times10^6$). Counts per million (CPM) were subsequently derived from library-size–normalized read counts [6, 7].

**Environmental sample processing (Barston STW)**

Raw sewage (post-grill) was collected at Barston Sewage Treatment Works (Birmingham, UK) on 9 October 2023, stored at 4 °C, and sequentially filtered (0.45 µm and 0.22 µm pore-sized filters) to remove particulates. The viral fraction was concentrated approximately 100-fold using centrifugal ultrafiltration (100 kDa MWCO). *E. coli* PB114 minicells ($2.1 \times 10^9$ cells/mL) were incubated with viral concentrates in three biological replicates. Following interaction and washing (as described above), DNA was extracted, purified, and subjected to whole-genome amplification, with negative controls included throughout.

**Library preparation and sequencing (environmental datasets)**

Library preparation and sequencing were performed at the Centre for Genomic Research/NERC Environmental Omics Facility. Libraries were prepared using the NEBNext Ultra II FS DNA Library Prep Kit with reduced reaction volumes and unique dual indexing, followed by limited PCR amplification. Libraries were purified, quantified, size-validated, pooled equimolarly, and sequenced on an Illumina NovaSeq X Plus platform (10B flow cell) to generate paired-end reads, corresponding to approximately 2.69 billion trimmed reads.

**Metagenomics workflow analysis (environmental datasets)**

Preprocessing (performed by NERC E/O Facility): Cutadapt v1.2.1 [8] with option -O 3 to trim adapter matches ≥3 bp at 3′ ends; Sickle v1.2 [9] with a minimum window quality score of 20; reads <15 bp removed; if only one read in a pair passed, it was retained in R0. Statistics from EAUtils fastq-stats [10]. Error correction was performed using BBMap's Tadpole module (v38.84; mode= correct, ecc= t, prefilter= 2) [30]. Read normalisation and deduplication were carried out using BBMap's Clumpify module (v38.84; groups= auto, dedupe= t, subs= 0, passes= 2) [11, 12]. Assembly: SPAdes v4.0 with -meta flag [13]. Assemblies were combined; contigs were renamed sequentially to vOTU_Sample_X (X starting at 1) using change_headers.py (Python 3.10.8; Supplementary Data 9). Redundancy removal: All vOTUs (three total viromes and three *E. coli* minicell-enriched viromes; six assemblies total, including replicates) were concatenated; a 10 kb cutoff was applied to remove short contigs [12].

BLASTn was used to compute Average Nucleotide Identity (ANI) and Alignment Fraction (AF). anicalc.py and aniclust.py clustered contigs at ≥ 95% ANI and ≥ 85% AF [14]. Cluster membership was written to my_clusters.tsv; unique identifiers were extracted with Final_vOTUs_unique_extract.py to produce the deduplicated FASTA (Supplementary Data 10). Viral identification/cleaning: geNomad v4.1.0 with option --clean-up [15] on the deduplicated vOTUs FASTA. Host sequence exclusion: Filtered viral vOTUs were aligned against host genomes with minimap2 v2.24-r1122 using options -a -x asm5 [5]; SAMtools v1.13 applying the -f 4 filter [16] to retain only unmapped reads. Quality assessment: CheckV v1.5.0 (default settings) [14] to remove host flanking regions, estimate completeness via AAI, assign MIUViG quality tiers (complete, high, medium, low, undetermined), and flag circular genomes. Annotation: Metaprokka v1.15.0 [15, 17, 18] using PHROGs HMM database v4 with default parameters. Host prediction: iPHoP v1.0.0 [19] (Aug_2023 database; default parameters) to assign genus-level hosts using integrated phage- and host-based modules. Subsampling: To harmonise depth across libraries, host-filtered trimmed reads were counted with SeqFu v1.20.3 (count) [4] to identify the minimum library size, and all samples were subsampled to this depth using seqtk v1.4 (r122). Mapping: BBMap v38.84 [3] with minid= 0.95 to map subsampled reads to (i) final vOTUs and (ii) host genomes. Abundance and filters: Sequenced reads of unenriched and enriched viromes were normalised using counts per million (CPM) [20, 21] and centered log-ratio (CLR), ensuring observed community differences reflect biology rather than sequencing depth or compositional effects [22]. A vOTU was retained if it had > 70% genome coverage and an average fold-change >1 across three biological replicates (union of replicates rule), to meet community standards [23].

A genome-wide proteomic tree was generated using ViPTree [24] based on normalized tBLASTx similarities between viral genomes and midpoint-rooted for visualization. TerL sequences from representative crass-like phages across the order *Crassvirales* (ICTV VMR; MSL40.v2) were aligned using MAFFT v7 (option --auto) [25]. A maximum-likelihood phylogeny was inferred using IQ-TREE2 with ModelFinder (-m MFP) [26] and ultrafast bootstrap and SH-aLRT support (1000 replicates) [27]. The tree was visualized using iTOL [28], with phage T7 (NC_001604) TerL as an outgroup Supplementary Tables 12 (08_crAss_like Phage_Tree). *E. coli* enriched vOTUs were compared against the complete phage genome database available as of 31 August 2025 [29] using Mash v2.3 with k-mer size 21 and sketch size 1000. The closest match per vOTU was retained based on minimum Mash distance. Estimated ANI was calculated as (1 −Mash distance) × 100.

**Visualisation and statistics:** Analyses in Python 3.12.4 with Pandas 2.2.2 and NumPy; statistics with SciPy [30], scikit-learn [31], statsmodels [32], scikit-bio[33]; plots with Plotly [34] and Matplotlib 3.9.1 [35]. Alpha-diversity (Chao1 and Shannon) computed on relative abundances; group differences by Kruskal-Wallis with Dunn's post-hoc, FDR-controlled. Beta-diversity: Bray-Curtis on square-root-transformed CPM and Jaccard on presence/absence, visualised by NMDS. Per-vOTU differential enrichment is summarised as $\log_2$ (enriched/total) after adding a pseudocount of $1 \times 10^{-2}$. Histogram bin widths were determined using the Freedman-Diaconis rule (Histogram.py; Supplementary Data 12). vOTUs were classified as enriched ($\Delta$CLR $\geq +5$), depleted ($\Delta$CLR $\leq -5$), or unchanged otherwise, corresponding to a conservative ±2SD threshold (~32-fold change). Mean–difference (MA) plots (Scatterplot.py; Supplementary Data 14) were used to visualise $\Delta$CLR versus mean CLR abundance, with ±5 $\log_2$ units applied as enrichment thresholds. For each library, the mapping fraction F to the *E. coli* PB114 (MG1655-derived) genome as a host-mapping fraction was calculated as $F = M/T$, where M is the aligned reads, and T is the total raw reads. Values reported per replicate in Supplementary Tables 12 (02_Abundance).

**Electron microscopy**

Purified *E. coli* PB114 minicells incubated with phage mixtures were pelleted at 16 000 × g for 5 min and fixed in 2.5% (v/v) glutaraldehyde in SM buffer at 4 °C. For transmission electron microscopy (TEM), samples were applied to glow-discharged carbon-coated copper grids, washed with water, negatively stained with uranyl acetate, and air-dried before imaging on a JEOL JEM-1400 microscope operated at 120 kV. For scanning electron microscopy (SEM), fixed minicells were deposited onto poly-L-lysine–coated glass slides, post-fixed with osmium tetroxide, dehydrated through an ethanol series, treated with hexamethyldisilazane, sputter-coated with platinum, and imaged using a Zeiss Gemini 360 microscope.

### Results

Supplementary Figure 1. Microscopic and quantitative assessment of minicell purification from *E. coli* PB114. (1A) SEM image showing purified minicells (~200-400 nm) predominantly clustered together, with minor aggregation, also confirming high visual purity. (1B) Comparative analysis of viable *E. coli* PB114 cells (red bars, CFU mL$^{-1}$) and total cells including minicells (blue bars, cells mL⁻¹) during purification based on $OD_{600}$ measurements. Steps: A = growth on LB agar plates after initial culture; B-D = differential centrifugation to isolate minicells; E-H = antibiotic treatment followed by additional centrifugation; I = filtration through a 0.7 µm syringe filter; J = final concentration and purity assessment. Total cell number corresponds to viable + minicell counts per mL.

1A

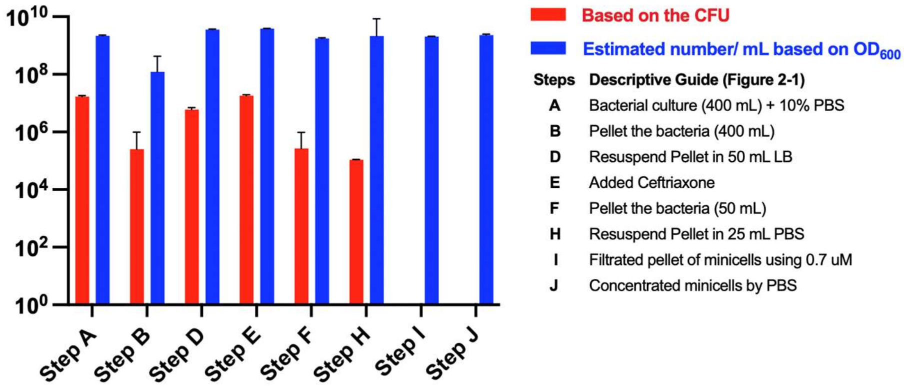


1B

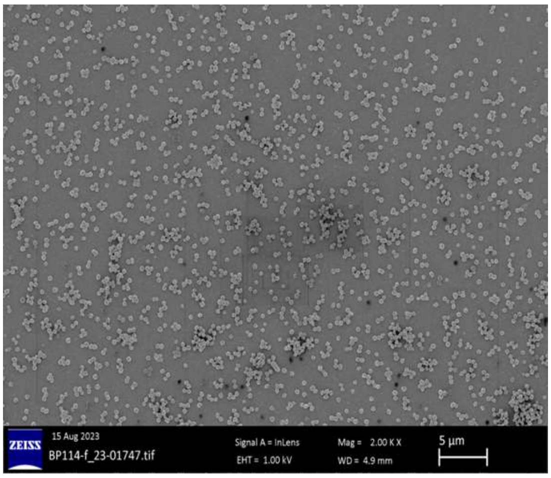


Supplementary Figure 12. Interaction of *E. coli* PB114 minicells with host-specific and non-host phages. (1A) Interaction of *E. coli* PB114 minicells with phages yaya_002 (*E. coli*-specific) and AA0033 (*S. capitis*-specific). The unenriched virome contained $2 \times 10^3$ PFU/mL of yaya_002 and $7.33 \times 10^8$ PFU/mL of AA0033 (ratio $1 : 3.7 \times 10^5$). After incubation with minicells, the yaya_002 titer decreased to $2 \times 10^2$ PFU/mL (90% reduction), while AA0033 remained unchanged at $8 \times 10^8$ PFU/mL, resulting in a post-enrichment ratio of $1 : 4 \times 10^6$. Error bars indicate standard deviations from replicate experiments. (1B) SEM of *E. coli* PB114 minicells interacting with the *E. coli*-specific phage yaya_002. Black arrows indicate phage particles; red arrows indicate *E. coli* PB114 minicells.

2A

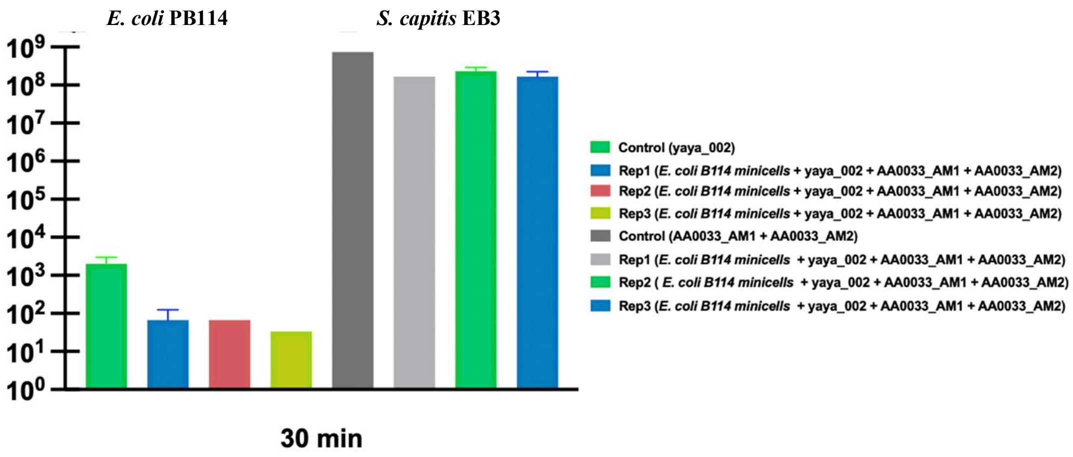


2B

2C

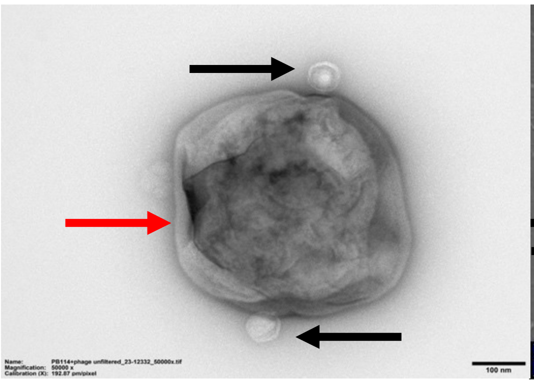


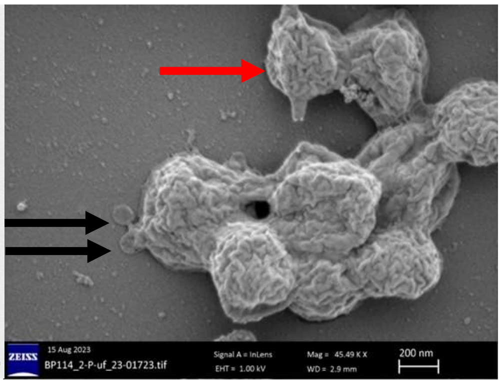


Supplementary Figure 2. (2A) Interaction of *E. coli* PB114 minicells with phages yaya_002 (*E. coli*–specific) and AA0033 (*S. capitis*-specific). The baseline virome contained 2 × $10^3$ PFU/mL of yaya_002 and 7.33 × $10^8$ PFU/mL of AA0033 (ratio 1:3.7 × $10^5$). After incubation with minicells, yaya_002 decreased to 2 × $10^2$ PFU/mL (90% reduction), while AA0033 remained unchanged at 8 × $10^8$ PFU/mL, yielding a post-enrichment ratio of 1:4 × $10^6$. *E. coli* PB114 and *S. capitis* served as lawn bacteria for plaque assays. Error bars represent standard deviations from biological triplicates. (2B) TEM imaging of *E. coli* PB114 minicells interacting with phage yaya_002. Black arrows indicate phage particles; red arrows indicate minicells. (2C) SEM imaging of *E. coli* PB114 minicells interacting with phage yaya_002. Black arrows indicate phage particles; red arrows indicate minicells.

Supplementary Table 2- Summary of read mapping to the *E. coli* MG1655 reference genome for proof of principle. The table lists total raw reads (T), aligned reads (M), and the corresponding mapping fraction (F = M/T).

| ID | Total Raw Reads (T) | Aligned Reads (M) | Mapping Fraction (F=M/T) % |
|---|---|---|---|
| **Control1** | 4.19E+06 | 2.56E+03 | 6.11E-02 |
| **Rep1** | 3.68E+06 | 6.13E+03 | 1.67E-01 |
| **Rep2** | 4.03E+06 | 7.5E+03 | 1.86E-01 |
| **Rep3** | 6.26E+06 | 1.18E+04 | 1.88E-01 |

Supplementary Table 3. List of Barston Sewage Treatment Works (STW) samples analyzed before and after minicell interaction. Fresh raw sewage collected on 9 October 2023 from Barston Sewage Treatment Works (Birmingham, UK) was concentrated to obtain environmental viromes, which were subsequently incubated with *E. coli* PB114 (*ΔminC*) minicells. Samples included total viromes, minicell-interacted viromes, and an *E. coli* PB114 minicell negative control, each processed in triplicate, yielding a total of nine samples. The table also includes the coliphage isolate APPB2_1 derived from the sewage sample. Raw and trimmed read counts, as well as corresponding run and study accession numbers deposited in the European Nucleotide Archive (ENA), are presented.

| Name | Description | Incubated with *E. coli* PB114 | Genome Length / Raw Reads Counts | Trimmed Reads Counts | Run Accession (ENA) | Study Accession (ENA) |
|---|---|---|---|---|---|---|
| **TVB1** | Total Virome Replicate 1 | No | 443193914 | 440793434 | ERR15368070 | PRJEB72100 |
| **TVB2** | Total Virome Replicate 2 | No | 500019774 | 497625418 | ERR15368071 | PRJEB72100 |
| **TVB3** | Total Virome Replicate 3 | No | 444478440 | 442879572 | ERR15365195 | PRJEB72100 |
| **MEPB1** | *E. coli* PB114-Enriched Virome Replicate 1 | Yes | 182294908 | 181799112 | ERR15368064 | PRJEB72100 |
| **MEPB2** | *E. coli* PB114-Enriched Virome Replicate 2 | Yes | 178936998 | 175579378 | ERR15368065 | PRJEB72100 |
| **MEPB3** | *E. coli* PB114-Enriched Virome Replicate 3 | Yes | 197615040 | 194919892 | ERR15368066 | PRJEB72100 |
| **NC_MEPB** | Only Minicells of *E. coli* PB114 (Negative control) | – | Failed | – | – | – |
| **APPB2_1** | Host and propagated: *E. coli* PB114 | No | 165933 bp | – | | |

Supplementary Table 4. Summary of read mapping to the *E. coli* MG1655 reference genome for total and minicell-enriched virome samples. Quality-filtered sequencing reads from total virome samples (TVB1-3) and *E. coli* PB114 minicell-enriched virome samples (MEPB1-3) were aligned to the *E. coli* MG1655 genome to assess potential host-genome contamination. The table lists total raw reads (T), aligned reads (M), and the corresponding mapping fraction (F = M/T). Low mapping fractions in both total (mean 0.23%) and minicell-enriched (mean 1.12%) samples indicate that host-derived DNA remained minimal following minicell interaction.

| ID | Total Raw Reads (T) | Aligned Reads (M) | Mapping Fraction (F=M/T) % |
|---|---|---|---|
| TVB_1 | 4.408E+08 | 1.2604E+04 | 0.29 |
| TVB_2 | 4.976E+08 | 1.0008E+04 | 0.20 |
| TVB_3 | 4.429E+08 | 1.0008E+04 | 0.23 |
| MEPB_1 | 1.818E+08 | 4.194E+03 | 0.23 |
| MEPB_2 | 1.756E+08 | 2.0069E+04 | 1.14 |
| MEPB_3 | 1.949E+08 | 3.8706E+04 | 1.99 |

**3A)** Alpha-Diversity-Chao1

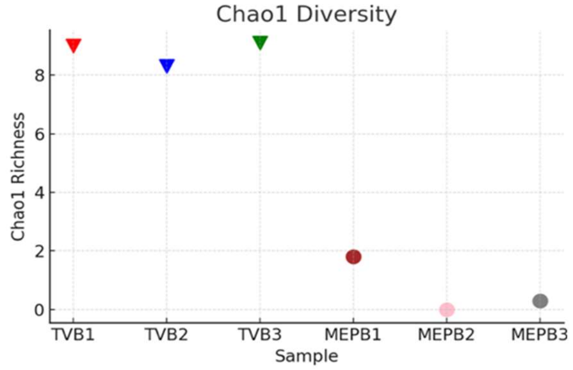


3B) Alpha-Diversity-Shannon

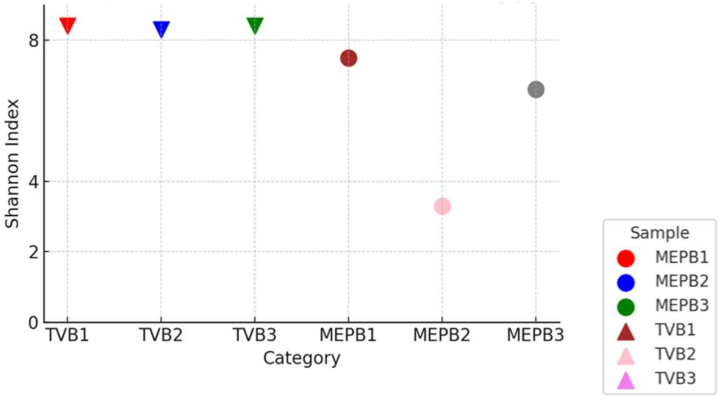


**3C) Beta-Diversity- Jaccard**

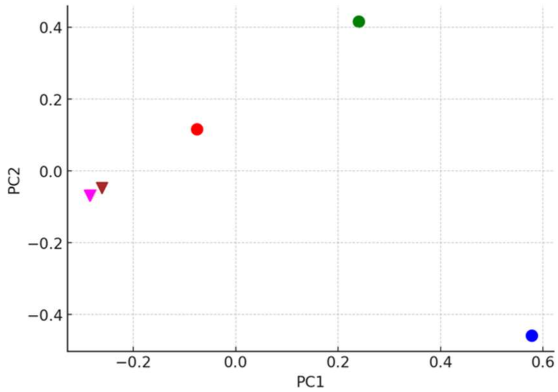


**3D) Beta-Diversity-Bary-Curtis**

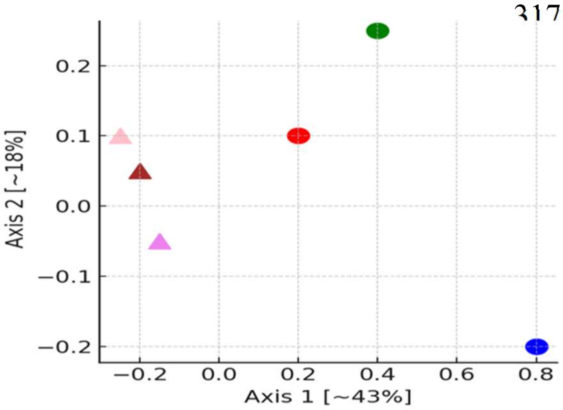


**4E) Alpha-Diversity**

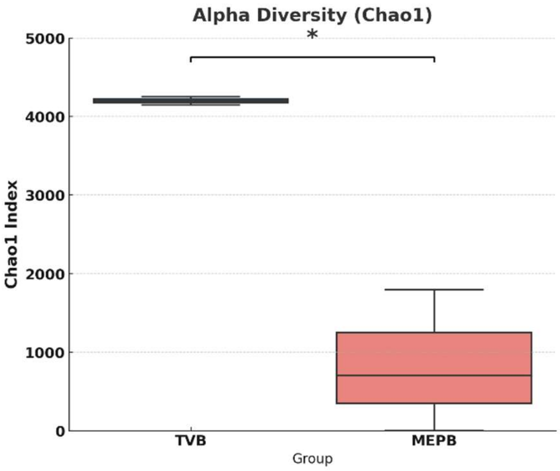


**4F) Beta-Diversity**

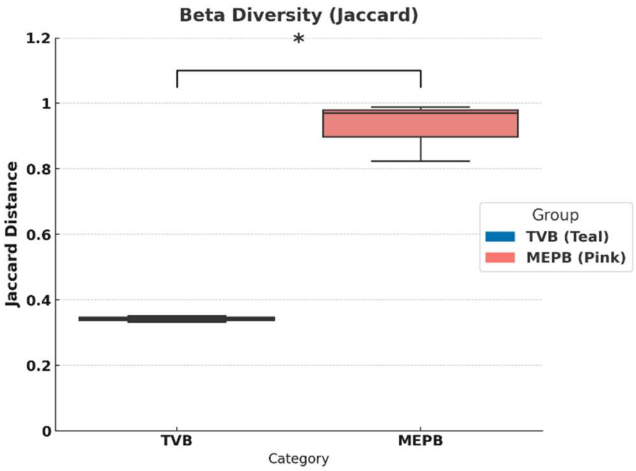


Supplementary Figure 3. Abundance and fold-Change of vOTUs in *E. coli* PB114-enriched vs. Total Viromes in metagenomics analysis. Figure 1A: Alpha diversity Chao 1 comparison of α-diversity metrics among virome replicates. Figure 1 B: Beta diversity Chao 1 comparison of B-diversity metrics among virome replicates. The experiment is conducted using biological replicates.

Supplementary Table 5. Kruskal-Wallis H-test results for Chao1 index comparisons among total virome (TVB) and *E. coli* PB114-enriched virome (MEPB) samples. Alpha-diversity based on the Chao1 richness index was compared among the three sample groups to evaluate the effect of minicell enrichment on viral community richness. The Kruskal-Wallis test indicated a significant difference between the total virome (TVB) and minicell-enriched virome (MEPB) samples (H = 3.971, P = 0.046), suggesting that minicell enrichment reduced overall virome richness. No significant differences were observed for other pairwise or overall group comparisons.

| Comparison | H Statistic | P-value |
|---|---|---|
| TVB vs MEPB | 3.971 | 0.046* |
| TVB vs MSTB | 0.441 | 0.507 |
| MEPB vs MSTB | 1.190 | 0.275 |
| All groups (TVB, MEPB, MSTB) | 3.854 | 0.146 |

*Significant at $P < 0.05$.

Supplementary Table 6. Dunn's post-hoc test for pairwise Chao1 index comparisons among virome sample groups. To identify which specific groups differed in alpha-diversity, Dunn's post-hoc test was applied following the Kruskal-Wallis analysis. The comparison confirmed a significant difference in Chao1 richness between MEPB and TVB samples (P = 0.0463), consistent with the observed reduction in diversity following minicell enrichment. Other pairwise contrasts were not significant.

| Comparison | Z | P.unadj | P.adj |
|---|---|---|---|
| MEPB - TVB | -1.9926 | 0.0463 | 0.0463* |

* Significant at $P < 0.05$.

Supplementary Table 7. Within-group variation in alpha-diversity (Chao1 index) between total virome and *E. coli* PB114-enriched virome (MEPB). Within-group variation was evaluated using standard deviation (SD) and two-standard-deviation (2SD) cut-offs to assess consistency among biological replicates. The relatively low SD observed in MEPB replicates (2.77; 2SD cutoff = 5.54) indicates that reductions in alpha-diversity were consistent across replicates.

| Label | sd_M1 | 2SD_cut-off |
|---|---|---|
| **MEPB** | 2.770776913 | 5.541553826 |

Supplementary Table 8. Summary statistics of ΔCLR values across all vOTUs used to define enrichment thresholds for histogram and mean–difference (MA) plot analyses. The global mean ($-2.6 \times 10^{-16}$) and standard deviation (SD = 2.71) were used to calculate ±2SD thresholds (±5.42 $\log_2$ units) for classification of enriched and depleted vOTUs.

| global_mean | global_sd | upper_mean_plus_2sd | lower_mean_minus_2sd |
|---|---|---|---|
| -2.57E-16 | 2.70865931 | 5.417318617 | -5.417318617 |

Supplementary Figure 4. Predicted phage host orders in the total virome (TVB) based on iPHoP analysis (v4.2.7.10). Host associations were inferred for 6181 viral operational taxonomic units (vOTUs) to establish baseline host range prior to minicell interaction. High-confidence (≥ 90%) genus-level predictions were obtained for 3145 vOTUs (51%), while 1996 vOTUs (64%) remained unassigned ("N/A"), reflecting substantial viral novelty. Among classified vOTUs, Bacteroidales was the dominant predicted host order (810 vOTUs; 27.3%), followed by Pseudomonadales (102; 3.4%), Oscillospirales (71; 2.4%), and Enterobacterales (64; 2.2%). This profile indicates a predominance of Bacteroidales-infecting phages in the sewage virome and provides a baseline for comparison with minicell-enriched fractions.

Supplementary Table 9. To evaluate the performance of the minicell enrichment platform, plaque assays were performed using *E. coli* PB114 as the host. Two phages infecting *E. coli* PB114 (APPB_002 and APPB_003) were isolated and genomically characterized. Sequencing reads from these isolates were mapped to both total (TVB) and *E. coli* PB114-enriched (MEPB) viromes and counts per million (CPM) were calculated for each condition. Mean CPM values increased markedly in the enriched virome, with APPB2_002 showing a 17.8-fold (1677%) increase relative to the total virome. These results confirm strong selective enrichment of *E. coli* PB114-specific phages following minicell-based enrichment.

| ID | CPM Mean TVB | CPM Mean MEPB | MEPB / TVB Ratio | Percent Change |
|---|---|---|---|---|
| **APPB2_1** | 1.215E+04 | 2.159E+05 | 1.777E+01 | 1677.22% |

Supplementary Table 10. Genome completeness assessment of 6181 vOTUs based on CheckV analysis.

| | **High** | **Medium** | **Low** | **Unknown** | **Total (n)** | **Total (%)** |
|---|---|---|---|---|---|---|
| Complete (100%) | 1220 | 193 | 80 | 1 | 1494 | 24.2 |
| High (90-100%) | 55 | 54 | 28 | 0 | 137 | 2.2 |
| Medium (50- 90%) | 169 | 231 | 117 | 1 | 518 | 8.4 |
| Low (<50%) | 1081 | 1677 | 1212 | 19 | 3989 | 64.5 |
| Not determined | 0 | 0 | 0 | 43 | 43 | 0.7 |
| **Total** | 2525 | 2155 | 1437 | 64 | 6181 | 100 |

Supplementary Table 11. Genome completeness assessment of 225 minicells *E. coli* PB114-enriched virome based on CheckV analysis.

| **Completeness** | **High** | **Medium** | **Low** | **Total (n)** | **Total (%)** |
|---|---|---|---|---|---|
| Complete (100%) | 47 | 4 | 2 | 53 | 23.6 |
| High (90-100%) | 2 | 1 | 0 | 3 | 1.3 |
| Medium (50- 90%) | 9 | 5 | 5 | 19 | 8.4 |
| Low (<50%) | 38 | 57 | 51 | 146 | 64.9 |
| Not determined | 0 | 0 | 0 | 4 | 1.8 |

5A)

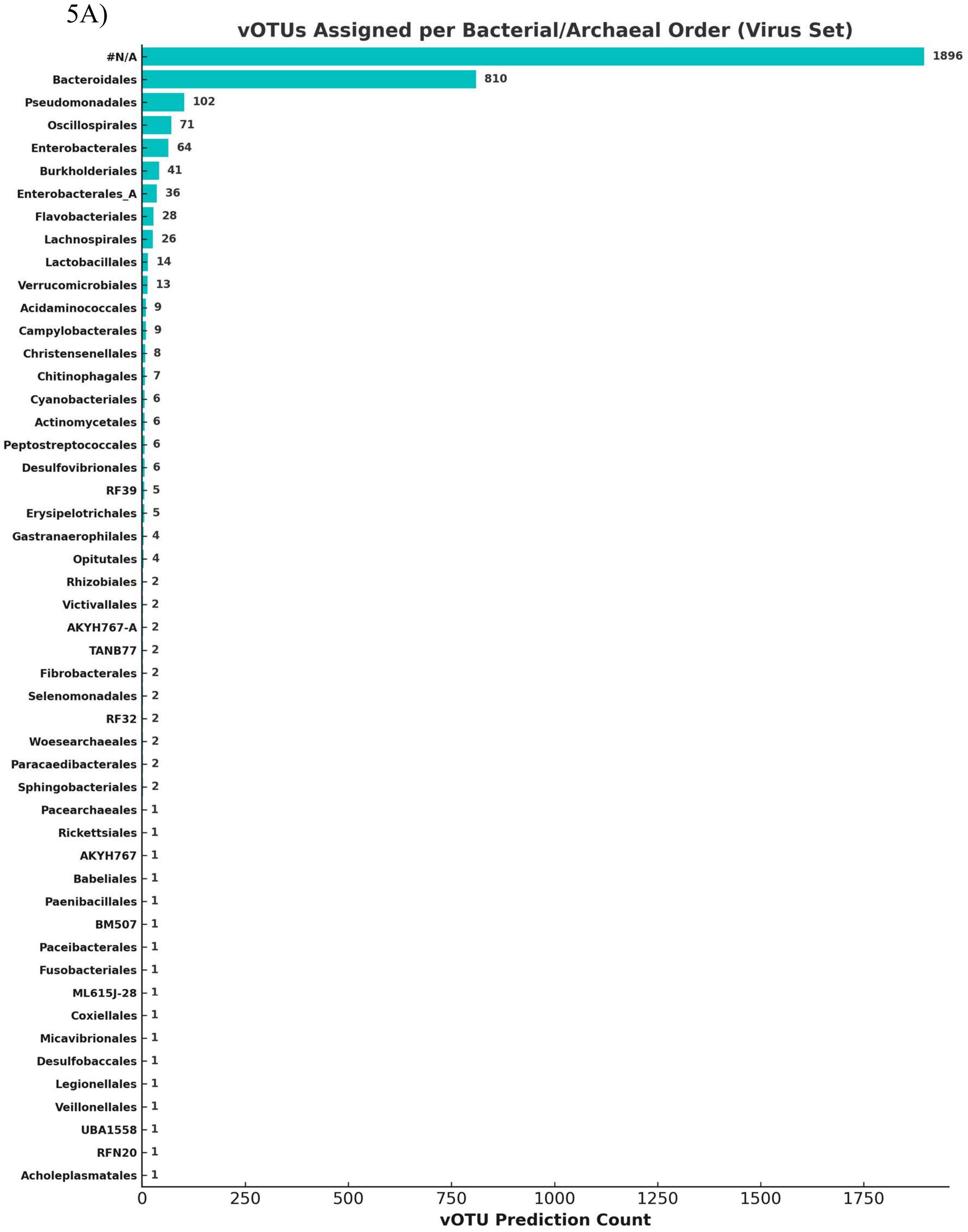
vOTUs Assigned per Bacterial/Archaeal Order (Virus Set)
#N/A 1896
Bacteroidales 810
Pseudomonadales 102
Oscillospirales 71
Enterobacterales 64
Burkholderiales 41
Enterobacterales_A 36
Flavobacteriales 28
Lachnospirales 26
Lactobacillales 14
Verrucomicrobiales 13
Acidaminococcales 9
Campylobacterales 9
Christensenellales 8
Chitinophagales 7
Cyanobacteriales 6
Actinomycetales 6
Peptostreptococcales 6
Desulfovibrionales 6
RF39 5
Erysipelotrichales 5
Gastranaerophilales 4
Opitutales 4
Rhizobiales 2
Victivallales 2
AKYH767-A 2
TANB77 2
Fibrobacterales 2
Selenomonadales 2
RF32 2
Woesearchaeales 2
Paracaedibacterales 2
Sphingobacteriales 2
Pacearchaeales 1
Rickettsiales 1
AKYH767 1
Babeliales 1
Paenibacillales 1
BM507 1
Paceibacterales 1
Fusobacteriales 1
ML615J-28 1
Coxiellales 1
Micavibrionales 1
Desulfobaccales 1
Legionellales 1
Veillonellales 1
UBA1558 1
RFN20 1
Acholeplasmatales 1
0
250
500
750
1000
1250
1500
1750
vOTU Prediction Count

5B)

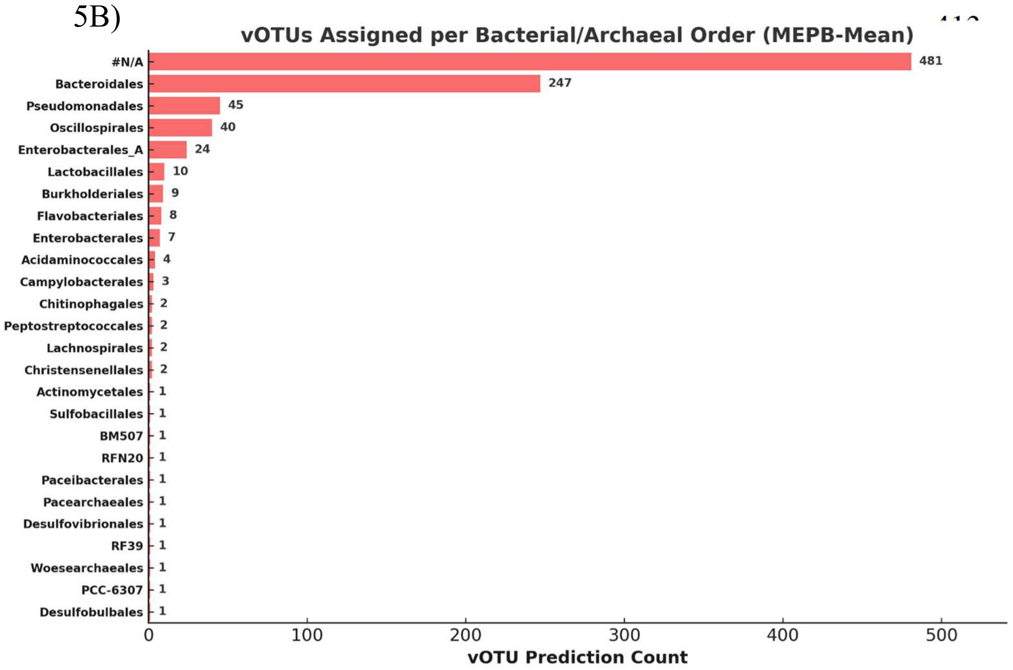


Supplementary Figure 5. A)Predicted phage host orders in the total virome (TVB) and B)*E. coli* PB114-enriched viromes (MEPB1-3) based on iPHoP (v4.2.7.10). Host predictions were compared across 6181 vOTUs to assess shifts following minicell enrichment. In the TVB, most vOTUs were unclassified ("N/A", 64.0%), with Bacteroidales (27.3%) as the dominant predicted host order. Following enrichment, the proportion of Enterobacterales-associated vOTUs increased (including Enterobacterales and Enterobacterales_A assignments), while unclassified vOTUs decreased to 59.8%. Enrichment ratios (MEPB/TVB $\geq 1$) indicate selective recovery of phages capable of interact with *E. coli* PB114 minicells.

Supplementary Table 12- Repository File Structure and Navigation Guide (Figshare Archive)

| Folder in Figshare | Description | Related Manuscript Section |
|---|---|---|
| **01_Metadata** | Proof: Metadata for proof-of-principle validation<br><br>Sewage: Metadata for sewage-derived total (TVB) and enriched (MEPB) viromes and Scripts | Methods – Proof-of-principle experiment (Supplementary Table 1)<br><br>Methods – Sewage-derived virome analysis (Supplementary Table 3) |
| **02_Abundance** | Proof: CPM calculations for defined phage mixture validation experiment; Host contamination<br>Sewage: CPM abundance matrix for total and enriched viromes; CLR-transformed abundance matrix and ΔCLR values; Host contamination | Proof-of-principle validation (Table 1-2)<br><br>CLR enrichment analysis (Fig. 1A-B; Supplementary Table 4) |
| **03_Quality_Check** | Viral genome classification summary; Genome completeness and contamination estimates | Genome quality assessment (Supplementary Table 10–11) |
| **04_Host_Prediction** | Host predictions (≥90% confidence) for enriched and total viromes | Host prediction analysis (Supplementary Figures 5–6) |
| **05_Mash** | Best Mash hit per enriched vOTU, including distance, ANI estimate, and shared hashes | Genome similarity analysis |
| **06_Coliphage_Isolate_Comparison** | Comparison of isolated coliphage APPB2_1 with enriched vOTUs | Supplementary Table 9 |
| **07_ViPTree** | Proteomic tree of 225 enriched vOTUs (Newick format) | Proteomic clustering analysis Fig. 1C |
| **08_crAss_like Phage_Tree** | | Fig. 1D |
| **09_ Statistics_results** | | |
| **10_Scripts** | | |

#### References


1. Andrews S Fastqc: A quality control tool for high throughput sequence data. Cambridge, United Kingdom.
2. Krueger F Trimgalore: A wrapper tool around cutadapt and fastqc to consistently apply quality and adapter trimming to fastq files. Babraham bioinformatics.
3. Bushnell B Bbmap. Sourceforge. Net/projects/bbmap. Accessed.
4. Telatin A, Fariselli P, Birolo G. Seqfu: A suite of utilities for the robust and reproducible manipulation of sequence files. *Bioengineering*. 2021;**8**:59
5. Li H. Minimap2: Pairwise alignment for nucleotide sequences. *Bioinformatics*. 2018;**34**:3094–100 https://doi.org/10.1093/bioinformatics/bty191
6. Ho SFS, Wheeler NE, Millard AD *et al.* Gauge your phage: Benchmarking of bacteriophage identification tools in metagenomic sequencing data. *Microbiome*. 2023;**11**:84 https://doi.org/10.1186/s40168-023-01533-x
7. Bikel S, López-Leal G, Cornejo-Granados F *et al.* Gut dsdna virome shows diversity and richness alterations associated to childhood obesity and metabolic syndrome. *iScience*. 2021;**24**:102900 https://doi.org/10.1016/j.isci.2021.102900
8. Martin M. Cutadapt removes adapter sequences from high-throughput sequencing reads. *EMBnetjournal*. 2011;**17**:10 https://doi.org/10.14806/ej.17.1.200
9. Joshi N, Fass J Sickle: A sliding-window, adaptive, quality-based trimming tool for fastq files (version 1.21)[software]; 2011.
10. Aronesty E. Ea-utils: Command-line tools for processing biological sequencing data. 2011
11. Bushnell B. Bbmap: A fast, accurate, splice-aware aligner. *United States*. Abstract Alignment of reads is one of the primary computational tasks in bioinformatics. Of paramount importance to resequencing, alignment is also crucial to other areas - quality control, scaffolding, string-graph assembly, homology detection, assembly evaluation, error-correction, expression quantification, and even as a tool to evaluate other tools. An optimal aligner would greatly improve virtually any sequencing process, but optimal alignment is prohibitively expensive for gigabases of data. Here, we will present BBMap [1], a fast splice-aware aligner for short and long reads. We will demonstrate that BBMap has superior speed, sensitivity, and specificity to alternative high-throughput aligners bowtie2 [2], bwa [3], smalt, [4] GSNAP [5], and BLASR [6]. AC02-05CH11231: Ernest Orlando Lawrence Berkeley National Laboratory, Berkeley, CA (US), 2014.
12. Roux S, Trubl G, Goudeau D *et al.* Optimizing de novo genome assembly from pcr-amplified metagenomes. *PeerJ*. 2019;**7**:e6902 https://doi.org/10.7717/peerj.6902
13. Bankevich A, Nurk S, Antipov D *et al.* Spades: A new genome assembly algorithm and its applications to single-cell sequencing. *Journal of Computational Biology*. 2012;**19**:455–77 https://doi.org/10.1089/cmb.2012.0021
14. Nayfach S, Camargo AP, Schulz F *et al.* Checkv assesses the quality and completeness of metagenome-assembled viral genomes. *Nature Biotechnology*. 2021;**39**:578–85 https://doi.org/10.1038/s41587-020-00774-7
15. Camargo AP, Roux S, Schulz F *et al.* Identification of mobile genetic elements with genomad. *Nature Biotechnology*. 2023 https://doi.org/10.1038/s41587-023-01953-y
16. Danecek P, Bonfield JK, Liddle J *et al.* Twelve years of samtools and bcftools. *GigaScience*. 2021;**10** https://doi.org/10.1093/gigascience/giab008
17. Seemann T. Prokka: Rapid prokaryotic genome annotation. *Bioinformatics*. 2014;**30**:2068–69 https://doi.org/10.1093/bioinformatics/btu153

18. Cook R, Brown N, Rihtman B *et al.* The long and short of it: Benchmarking viromics using illumina, nanopore and pacbio sequencing technologies. *Microbial Genomics*. 2024;**10**:001198
19. Roux S, Camargo AP, Coutinho FH *et al.* Iphop: An integrated machine learning framework to maximize host prediction for metagenome-derived viruses of archaea and bacteria. *PLOS Biology*. 2023;**21**:e3002083 https://doi.org/10.1371/journal.pbio.3002083
20. Leech J, Cabrera-Rubio R, Walsh AM *et al.* Fermented-food metagenomics reveals substrate-associated differences in taxonomy and health-associated and antibiotic resistance determinants. *mSystems*. 2020;**5** https://doi.org/10.1128/msystems.00522-20
21. McMurdie PJ, Holmes S. Waste not, want not: Why rarefying microbiome data is inadmissible. *PLoS Computational Biology*. 2014;**10**:e1003531 https://doi.org/10.1371/journal.pcbi.1003531
22. Gloor GB, Macklaim JM, Pawlowsky-Glahn V *et al.* Microbiome datasets are compositional: And this is not optional. *Frontiers in Microbiology*. 2017;**8** https://doi.org/10.3389/fmicb.2017.02224
23. Roux S, Adriaenssens EM, Dutilh BE *et al.* Minimum information about an uncultivated virus genome (miuvig). *Nature Biotechnology*. 2019;**37**:29–37 https://doi.org/10.1038/nbt.4306
24. Nishimura Y, Yoshida T, Kuronishi M *et al.* Viptree: The viral proteomic tree server. *Bioinformatics*. 2017;**33**:2379–80 https://doi.org/10.1093/bioinformatics/btx157
25. Katoh K, Standley DM. Mafft multiple sequence alignment software version 7: Improvements in performance and usability. *Molecular Biology and Evolution*. 2013;**30**:772–80 https://doi.org/10.1093/molbev/mst010
26. Minh BQ, Schmidt HA, Chernomor O *et al.* Iq-tree 2: New models and efficient methods for phylogenetic inference in the genomic era. *Molecular Biology and Evolution*. 2020;**37**:1530–34 https://doi.org/10.1093/molbev/msaa015
27. Hoang DT, Chernomor O, von Haeseler A *et al.* Ufboot2: Improving the ultrafast bootstrap approximation. *Molecular Biology and Evolution*. 2017;**35**:518–22 https://doi.org/10.1093/molbev/msx281
28. Letunic I, Bork P. Interactive tree of life (itol) v6: Recent updates to the phylogenetic tree display and annotation tool. *Nucleic Acids Research*. 2024;**52**:W78–W82 https://doi.org/10.1093/nar/gkae268
29. Cook R, Brown N, Redgwell T *et al.* Infrastructure for a phage reference database: Identification of large-scale biases in the current collection of cultured phage genomes. *PHAGE*. 2021;**2**:214–23 https://doi.org/10.1089/phage.2021.0007
30. McKinney W Data structures for statistical computing in python. *SciPy*. 51–56.
31. Pedregosa F, Varoquaux G, Gramfort A *et al.* Scikit-learn: Machine learning in python. *the Journal of machine Learning research*. 2011;**12**:2825–30
32. Seabold S, Perktold J. Statsmodels: Econometric and statistical modeling with python. In: *9th Python in Science Conference*, 2010.
33. Author. Scikit-bio/scikit-bio: Scikit-bio 0.7.1.Post1 [Computer software]. https://github.com/scikit-bio/scikit-bio/tree/0.7.1.post1. 2025.
34. Author. Plotly: Python graphing library [Computer software]. https://plotly.com/python/. 2015.
35. Hunter JD. Matplotlib: A 2d graphics environment. *Computing in science & engineering*. 2007;**9**:90–95
